# Cellular identity and Ca^2+^ signaling activity of the non-reproductive GnRH system in the *Ciona* larva

**DOI:** 10.1101/2020.01.02.893354

**Authors:** Nanako Okawa, Kotaro Shimai, Kohei Ohnishi, Masamichi Ohkura, Junichi Nakai, Takeo Horie, Atsushi Kuhara, Takehiro G. Kusakabe

## Abstract

Tunicate larvae have a non-reproductive gonadotropin-releasing hormone (GnRH) system with multiple ligands and receptor heterodimerization enabling complex regulation. In the *Ciona* larva, one of the *gnrh* genes, *gnrh2*, is conspicuously expressed in the motor ganglion and nerve cord, which are homologous structures to the hindbrain and spinal cord, respectively, of vertebrates. The *gnrh2* gene is also expressed in the proto-placodal sensory neurons, which are the proposed homologue of vertebrate olfactory neurons. The tunicate larvae occupy a non-reproductive dispersal stage, yet the role of their GnRH system remains elusive. In this study, we investigated neuronal types of *gnrh2*-expressing cells in *Ciona* larvae and visualized the activity of these cells by fluorescence imaging using a calcium sensor protein. Some cholinergic neurons and dopaminergic cells express *gnrh2*, suggesting that GnRH plays a role in controlling swimming behavior. However, none of the *gnrh2*-expressing cells overlap with glycinergic or GABAergic neurons. A role in motor control is also suggested by the correlation between the activity of some *gnrh2*-expressing cells and tail movements. Interestingly, *gnrh2*-positive ependymal cells in the nerve cord, known as a kind of glia cells, actively produced Ca^2+^ transients, suggesting that neuroendocrine signaling occurs in the glia cells of the nerve cord.

## Introduction

Gonadotropin-releasing hormone (GnRH) is a key regulator of reproductive functions in vertebrates^1,2^. GnRH has been suggested to play non-reproductive roles in the nervous system and during development^3–9^. Compared to its reproductive roles, the non-reproductive roles of GnRH are less well understood.

Tunicates are the sister group of vertebrates^10,11^. A conspicuous non-reproductive GnRH system has been reported in the larva of the sessile tunicate *Ciona*^12,13^. Six GnRH peptides and four receptors are encoded by the *Ciona* genome^14–18^. In the *Ciona* larva, the GnRH genes are strikingly expressed in the central nervous system (CNS) through the entire antero-posterior body axis^12^. Correspondingly, the GnRH receptor genes are specifically expressed in the tissues and organs located along the CNS, namely the notochord, the tail muscle, and the epidermal sensory neurons^12^. One of the *Ciona gnrh* genes, *gnrh2*, is conspicuously expressed in the motor ganglion and nerve cord of the larva, which are homologous structures to the hindbrain and spinal cord, respectively, of vertebrates. The *gnrh2* gene is also expressed in the proto-placodal sensory neurons, which are the proposed homologue of vertebrate olfactory neurons^19^. *Ciona* GnRH has been suggested to play a pivotal role in the control of metamorphosis^13^. Considering the complex and well-developed nature of the larval GnRH system in *Ciona*, GnRH may play diverse and important roles in developmental and physiological processes in *Ciona* larvae. To date, however, the roles of the *Ciona* GnRH system remain elusive.

In this study, we investigated neuronal types of *gnrh2*-expressing cells in the *Ciona* larva and visualized the activity of these cells by fluorescence imaging using a calcium sensor protein. Some cholinergic motor neurons as well as unique cholinergic cells along the nerve cord were found to express *gnrh2*, suggesting that GnRH plays a role in the control of swimming behavior. By contrast, none of the *gnrh2*-expressing cells overlapped with glycinergic or GABAergic neurons. A role in motor control was also suggested by the correlation between the activity of some *gnrh2*-expressing cells and tail movements. Interestingly, *gnrh2*-positive ependymal cells in the nerve cord, known as a kind of glia cells, produced Ca^2+^ transients, suggesting that active signaling, presumably involving GnRH, occurs in the nerve cord.

## Results

### *Gnrh2* is expressed in proto placode-derived sensory neurons and caudal glial ependymal cells

The 4.3-kb upstream region of *gnrh2* connected with a fluorescence reporter can recapitulate the expression patterns of *gnrh2*^12^ (Fig. 1). This upstream region was used to express mCherry and G-CaMP8 in cells expressing *gnrh2* (Figs. 1, 2 and 3). Cell types were identified by double fluorescent staining of larvae with cell type-specific markers.

**Figure 1.**
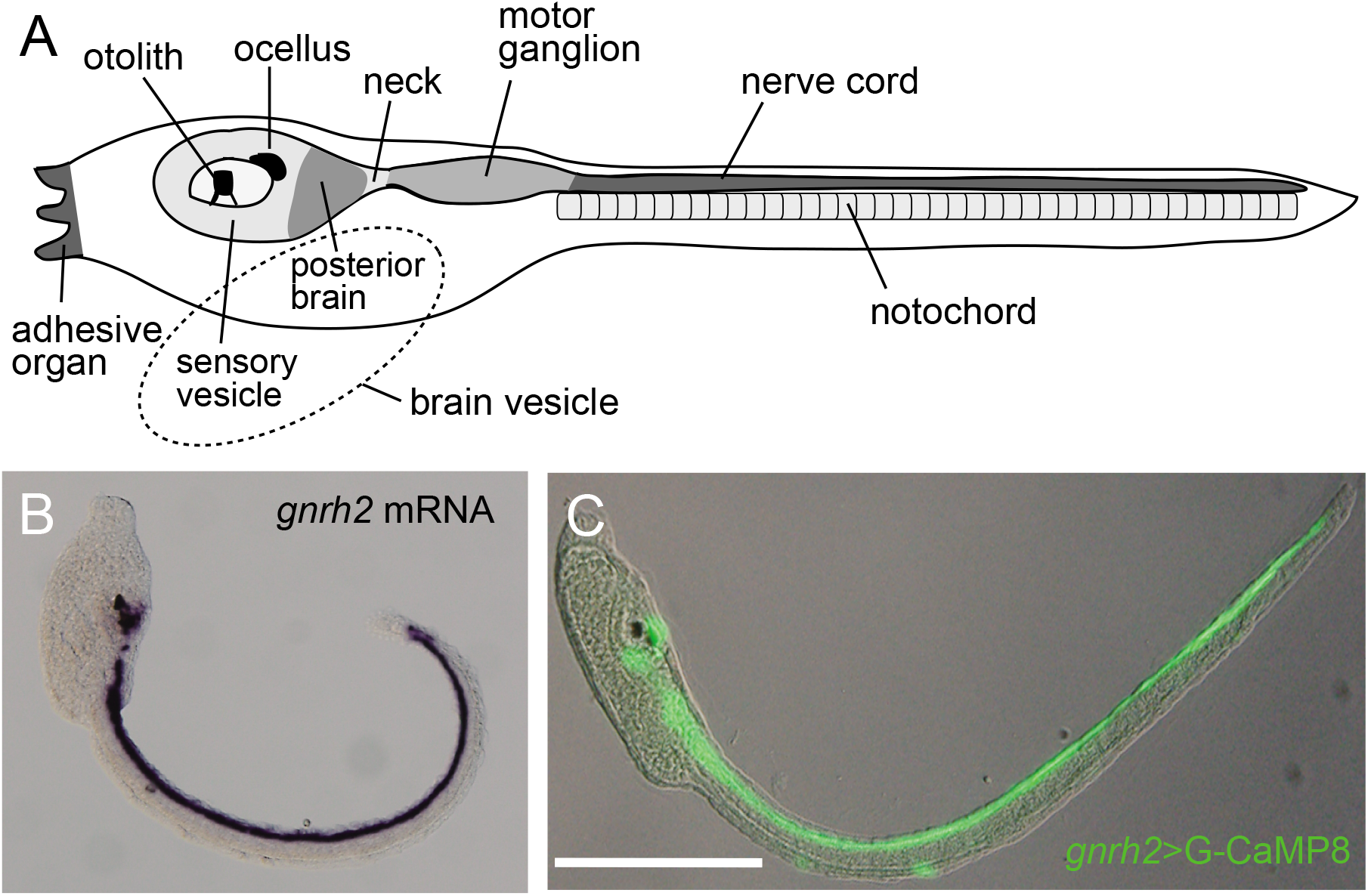
Expression patterns of *gnrh2* in the *Ciona* larva. (A) Schematic diagram showing the CNS of the larva. (B) Localization of *gnrh2* mRNA visualized by *in situ* hybridization. (C) Immunofluorescent localization of G-CaMP8 expressed under the control of the *cis*-regulatory region of *gnrh2.* Scale bar, 200 μm.

**Figure 2.**
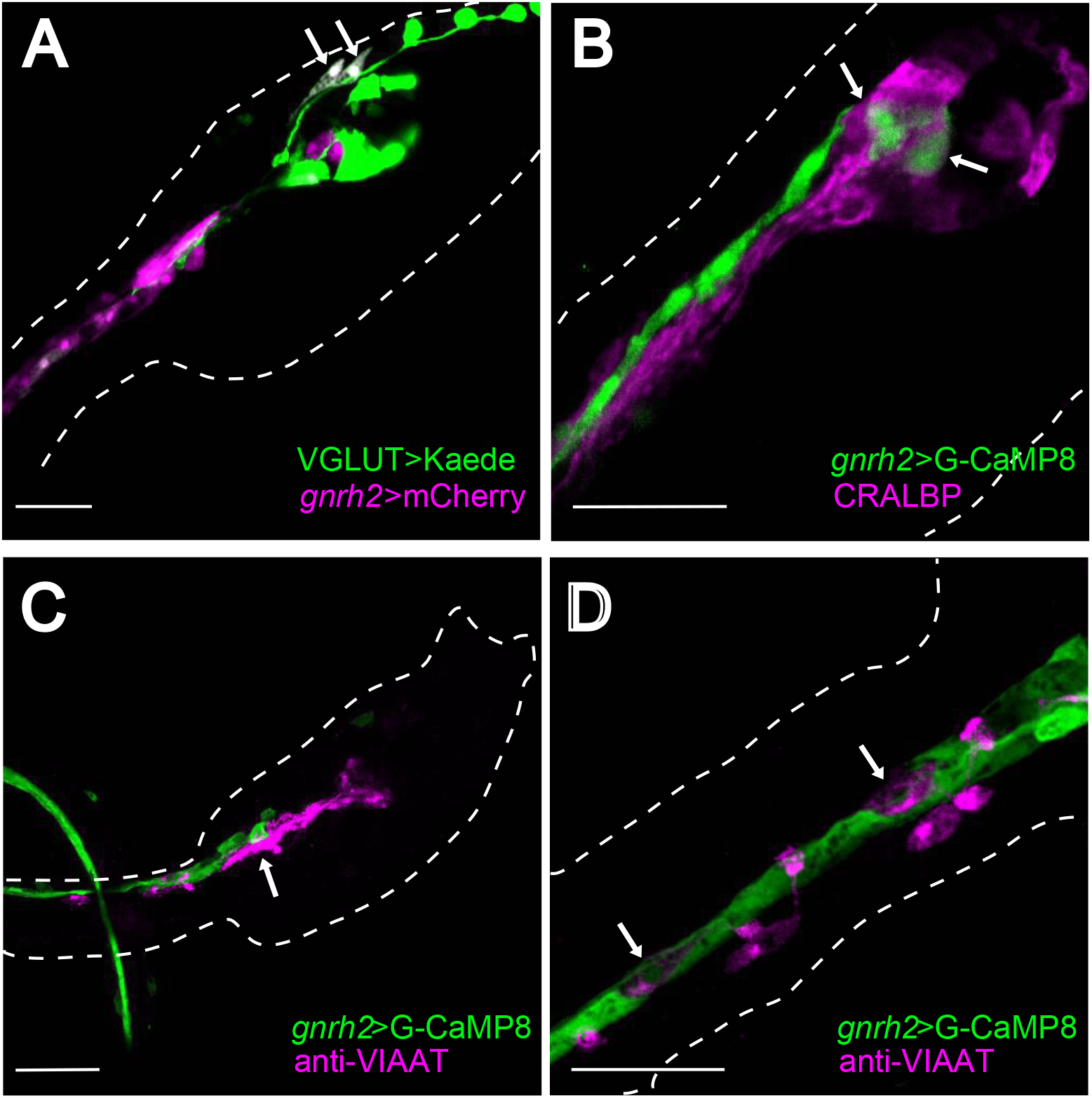
Immunohistochemical identification of types of cells expressing *gnrh2* in the *Ciona* larva. (A) Glutamatergic neurons and *gnrh2*-expressing cells were labeled with Kaede (*green*) and mCherry (*magenta*), respectively. The proto placode-derived sensory neurons (*arrows*) were shown to express *gnrh2*. (B) CRALBP-positive cells (*magenta*) were not overlapped with *gnrh2*-expressing cells (*green*). Arrows indicate *gnrh2*-expressing cells in the brain vesicle. (C,D) GABAergic/glycinergic neurons were visualized by immunostaining with anti-VIAAT antibody (*magenta*). *Arrows* in (C) indicate GABAergic/glycinergic neurons in the motor ganglion. *Arrows* in (D) indicate VIAAT-positive ACINs. Scale bars, 30 μm.

**Figure 3.**
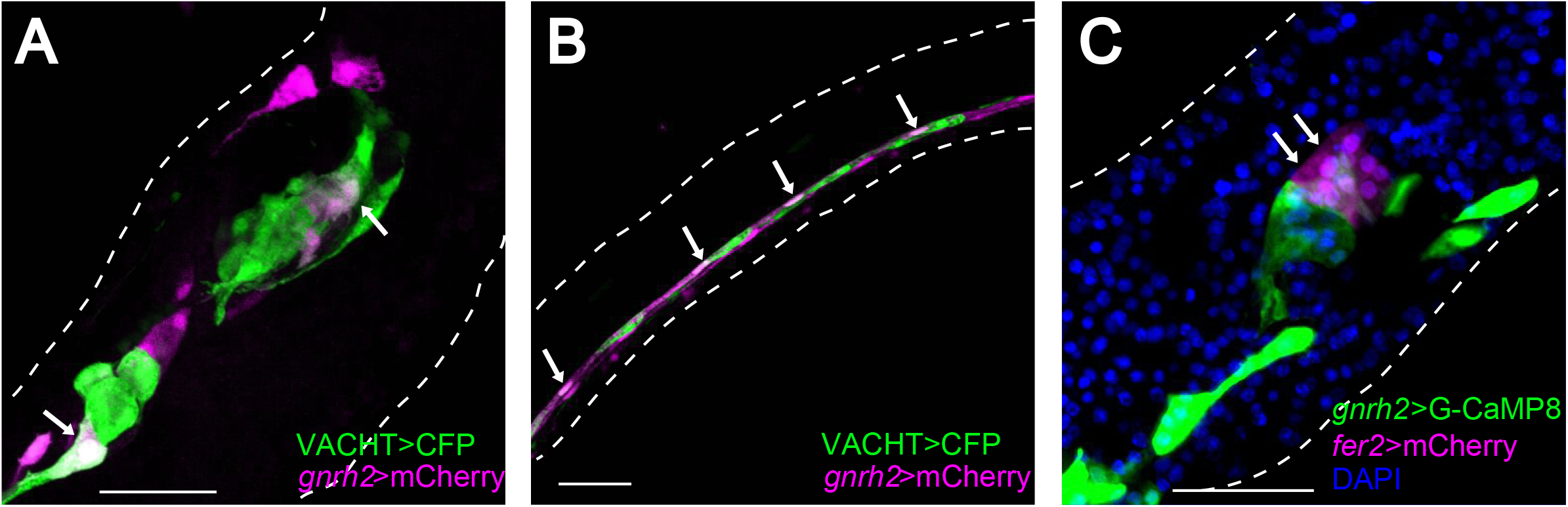
Some cholinergic and dopaminergic neurons express *gnrh2*. (A,B) Cholinergic neurons and *gnrh2*-exressing cells were labeled with CFP (*green*) and mCherry (*magenta*), respectively. *Arrows* indicate cells that co-expressed both markers. (C) Dopaminergic cells and *gnrh2*-expressing cells were labeled with mCherry (*magenta*) and G-CaMP8 (*green*), respectively. Some dopaminergic cells were also labeled with G-CaMP8 (*arrows*). Scale bars, 30 μm.

First, we examined whether *gnrh2*-expressing cells include glutamatergic neurons. In the *Ciona* larva, glutamate is a major neurotransmitter in the peripheral sensory neurons and photoreceptor cells^20^. Some interneurons in the posterior brain are also glutamatergic^20^. As previously reported^19^, proto-placode-derived *gnrh2*-expressing epidermal neurons (aATENs) are glutamatergic (Fig. 2A). aATENs are the only glutamatergic neurons that express *gnrh2*. None of the glutamatergic neurons express *gnrh2* in the CNS.

Next, we examined whether any of the GABAergic/glycinergic neurons express *gnrh2* using vesicular inhibitory amino acid transporter (VIAAT) as a marker. None of the VIAAT-positive cells overlapped with the reporter expression under the control of the *gnrh2 cis*-regulatory region (Figs. 2C and 2D). In the anterior tail region, there are two pairs of VIAAT-positive neurons called anterior caudal inhibitory neurons (ACINs), which align with glial ependymal cells in the lateral wall of the anterior nerve cord^21,22^. We found that the ACINs do not express *gnrh2*, whereas the lateral ependymal cells express *gnrh2* (Fig. 2D). The *gnrh2* expression in the lateral wall ependymal cells of the nerve cord is consistent with the *in situ* hybridization data previously reported^12^.

Cellular retinaldehyde-binding protein (CRALBP) is specifically localized in the glial ependymal cells in the brain vesicle and the motor ganglion^23,24^. In our immunohistochemicah analysis, CRALBP-positive cells were never overlapped with *gnrh2*-epxressing cells (Fig. 2B). Thus, in contrast to the conspicuous *gnrh2* expression in the ependymal cells of the nerve cord, *gnrh2* is not expressed in the ependymal cells in the brain vesicle and the motor ganglion.

### Some cholinergic and dopaminergic neurons express *gnrh2*

Acetylcholine is a major neurotransmitter at the neuromuscular junctions of the *Ciona* larva^21,25^. Cholinergic neurons were visualized by a fluorescence protein expressed under the control of the *cis*-regulatory region of the *vacht* gene^25^. *Gnrh2*-expressing neurons were shown to be cholinergic both in the brain vesicle and the motor ganglion (Fig. 3A).

The caudal part of the CNS (nerve cord) mainly consists of non-neuronal ependymal cells^26^. The nerve cord also contains two types of neurons: ACINs and bilateral pairs of cholinergic caudal neurons^21^. Some of these cholinergic caudal neurons seem to express *gnrh2* (Fig. 3B).

Another neurotransmitter that controls the swimming of *Ciona* tadpoles is dopamine^27^. Dopaminergic neurons are present in the brain vesicle^27–29^. Dopaminergic neurons were labeled with mCherry expressed under the control of the *cis*-regulatory region of the dopaminergic cell-specific gene *fer2*^27,29^ (previously described as *Ptf1a*, see Gyoja & Satoh^30^ for the orthologous families of the bHLH transcription factors). Double fluorescence imaging of dopaminergic neurons and *gnrh2*-expressing cells revealed that some dopaminergic neurons express *gnrh2* (Fig. 3C).

### Active Ca^2+^ transients in aATENs and *gnrh2*-expressing cells of the posterior CNS

G-CaMP8^31^ was used to monitor temporal changes in intracellular Ca^2+^ in *gnrh2*-expressing cells. Active Ca^2+^ transients were observed in aATENs, the motor ganglion, and the caudal nerve cord (Figs. 4-6; Movies S1-3). The larva contains two aATENs, and each has a sensory cilium^19^ (Fig. 4A,B). Both aATENs showed Ca^2+^ transients but their activities was were not synchronized (Fig. 4C; Movie S1).

**Figure 4.**
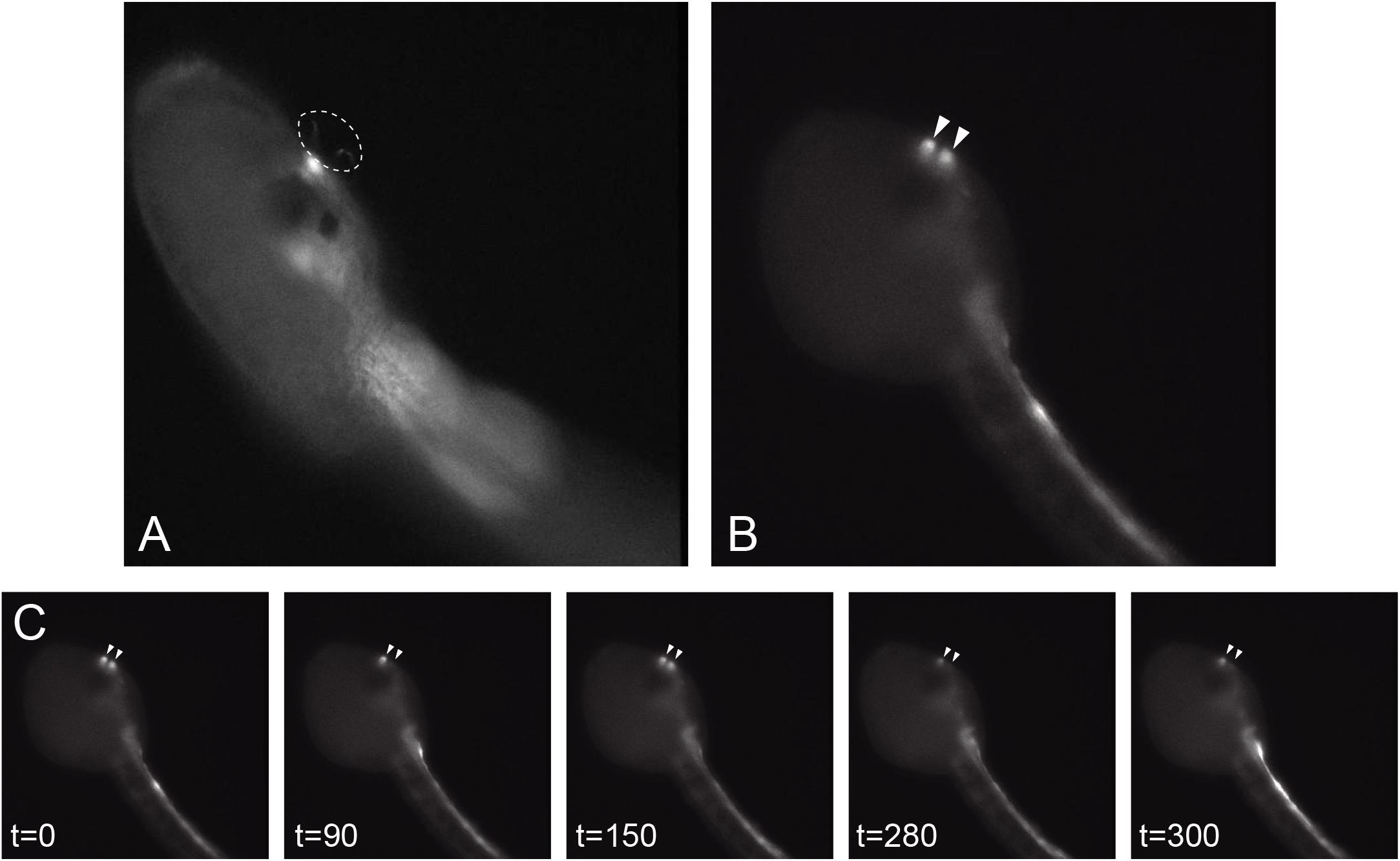
Calcium imaging of *gnrh2*-expressing chemosensory neurons. Fluorescence images of *Ciona* larvae expressing G-CaMP8 in the chemosensory aATENs. (A) The putative sensory cilia of each aATEN was labeled with G-CaMP8 fluorescence (dotted circle). (B,C) An example of a larva showing dynamic Ca^2+^ transients in the pair of aATENs (*arrowheads*) at 19 hours post fertilization (hpf). (C) Representative images of the larva recorded at the time (s) indicated. The serial images of the larva shown in (B,C) are shown in **Movie S1**.

Cholinergic *gnrh2*-expressing neurons at the posterior part of the motor ganglion exhibited active Ca^2+^ transients (Figs. 5 and 6). Periodic Ca^2+^ transients of a *gnrh2*-expressing neuron in the motor ganglion were observed at 19 hours after fertilization (*arrowheads* in Fig. 6B). In the tail region, Ca^2+^ transients were observed though the entire length of the nerve cord (Figs. 5 and 6). Both cholinergic neurons and ependymal cells showed Ca^2+^ transients in the tail nerve cord. For example, the narrow cell indicated by an *arrow* in Fig. 5A is presumably a cholinergic neuron. Many cells showing Ca^2+^ transients were block-shaped, which is characteristic of caudal ependymal cells (Fig. 6C).

**Figure 5.**
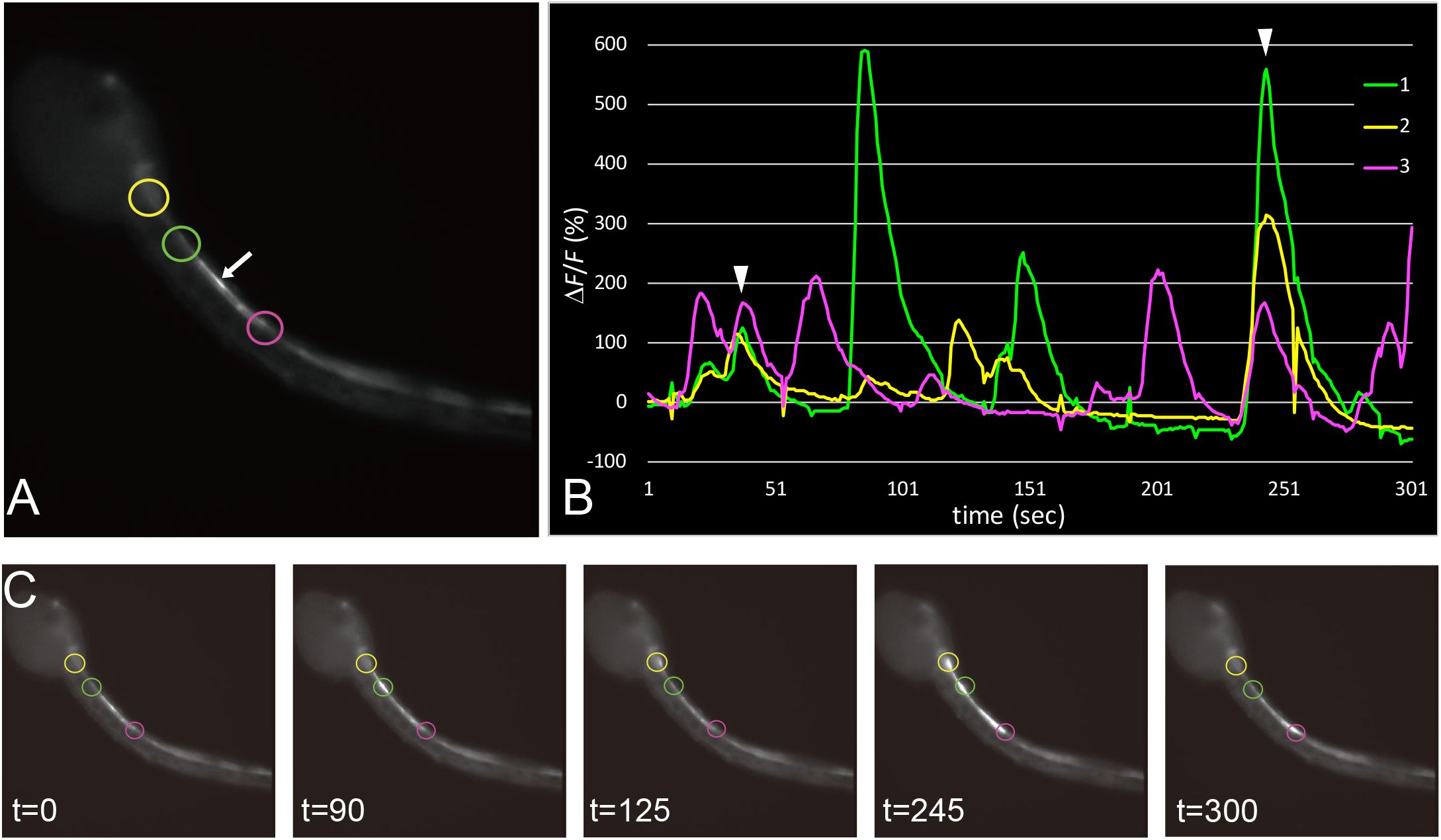
Interconnection between *gnrh2*-expressing cells in the larval CNS. (A) Fluorescence image of a larva at 19.5 hpf, showing G-CaMP8 fluorescence in the motor ganglion and the anterior nerve cord. (B) The graph shows the temporal patterns of fluorescence intensity at the three sites indicated by circles in (A). The colors of the lines correspond to the sites indicated by circles in the respective colors. The Ca^2+^ transients occurred independently of each other, but sometimes occurred at the same time (*arrowheads*). (C) Representative images of the larva recorded at the time (s) indicated. The serial images of the larva shown in (A) are shown in **Movie S2**.

**Figure 6.**
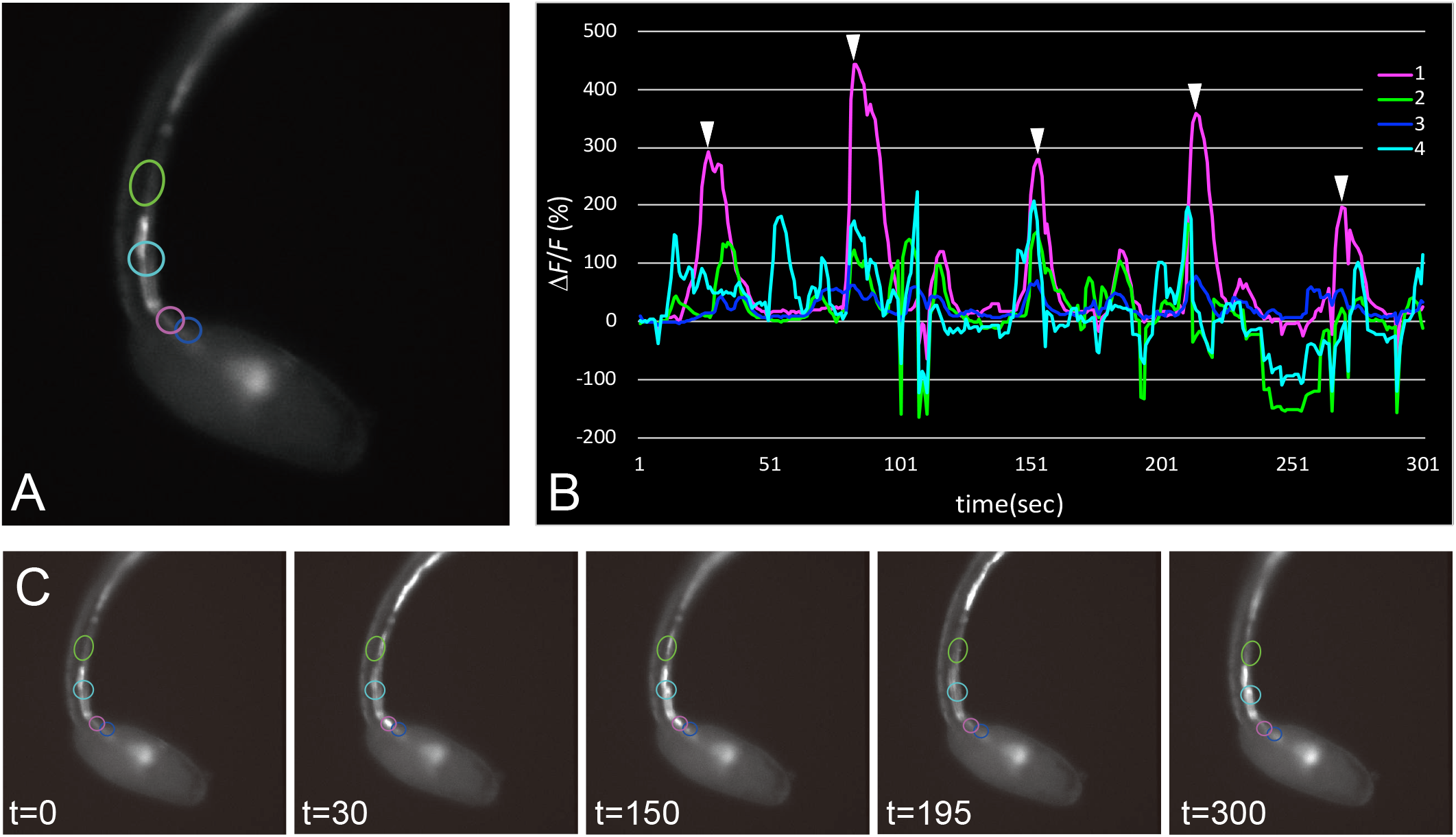
Periodic oscillation of Ca^2+^ transients in a *gnrh2*-expressing cell in the motor ganglion. (A) Fluorescence image of a larva at 20 hpf, showing G-CaMP8 fluorescence in the motor ganglion and the anterior nerve cord. (B) The graph shows the temporal patterns of fluorescence intensity at the four sites indicated by circles in (A). The colors of the lines correspond to the sites indicated by circles in the respective colors. Ca^2+^ spikes were periodically observed at regular intervals in the cell indicated by the magenta circle in (A) (*arrowheads*). (C) Representative images of the larva recorded at the time (s) indicated. The serial images of the larva shown in (A) are shown in **Movie S3**.

Ca^2+^ transients were not synchronized between cells— including neurons and ependymal cells—located at different sites of the larva (Figs. 5 and 6). However, simultaneous activation of cells at different sites was occasionally observed, as indicated by arrowheads in Fig. 5B and Fig. 6B, suggesting the presence of a neural circuit connecting *gnrh2*-expressing cells at different sites.

### Correlation between tail movements and Ca^2+^ transients in the motor ganglion and the anterior nerve cord

A neural circuit in the motor ganglion and the anterior nerve cord is thought to control muscle contraction in the tail^21,32,33^. Because active Ca^2+^ transients in *gnrh2*-expressing cells were observed in these regions, we examined the temporal correlation between tail movement and the activity of *gnrh2*-expressing cells. Ca^2+^ transients were frequently observed when the tail ceased its movement, whereas the Ca^2+^ transients were seemingly suppressed while the tail was moving (Fig. 7). Thus the tail movement often precedes the Ca^2+^ spike. This finding suggests the possible involvement of *gnrh2*-expressing cells in the control of swimming behavior.

**Figure 7.**
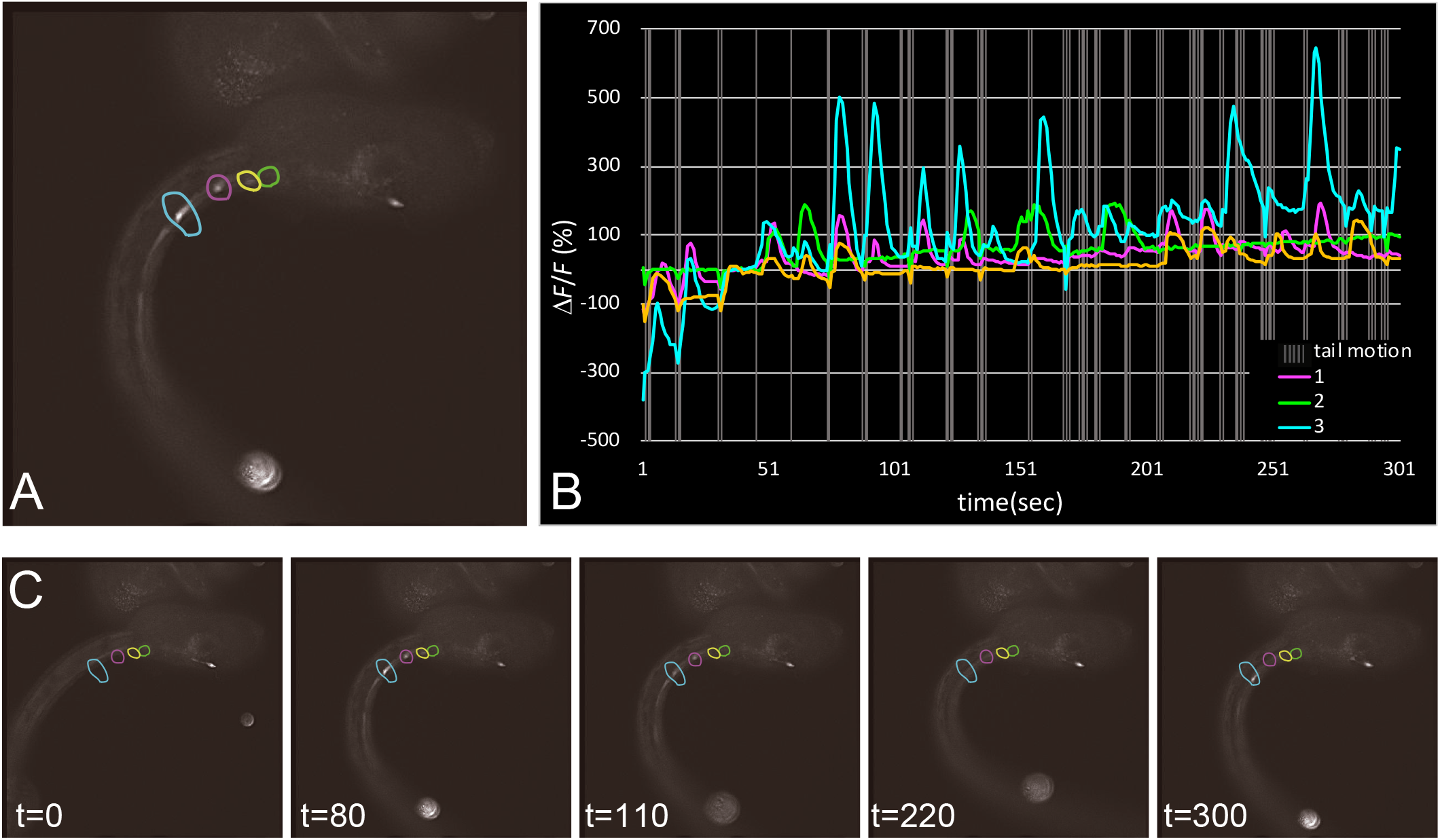
Correlation between Ca^2+^ transients in *gnrh2*-expressing cells and tail movement. (A) Fluorescence image of a larva at 19.5 hpf, showing G-CaMP8 fluorescence in the anterior nerve cord. (B) The graph shows the temporal patterns of fluorescence intensity at the four sites encircled by colored lines. The colors of the lines in the graph correspond to the sites encircled by lines in the respective colors. Gray vertical lines indicate the period when the tail was moving. Ca^2+^ transients generally occurred when the tail movement stopped. (C) Representative images of the larva recorded at the time (s) indicated. The serial images of the larva shown in (A) are shown in **Movie S4**.

## Discussion

In this study, we identified cell types of *gnrh2*-expressing cells and visualized their activity in the *Ciona* larva. Previously, the cells expressing GnRH-encoding genes had been only partially identified in *Ciona*. The caudal ependymal cells and the aATENs were reported to express *gnrh2*^12,19^. We confirmed these findings and further identified CNS neurons expressing *gnrh2*.

In the brain vesicle, dopaminergic neurons and a limited number of cholinergic neurons express *gnrh2*. Pharmacological and behavioral analyses have suggested that dopaminergic cells modulate the light-off-induced swimming behavior of *Ciona* larvae^27^. The role of cholinergic neurons in the *Ciona* brain vesicle has not been elucidated. Our present observations are the first to show the heterogeneity of cholinergic neurons in the brain vesicle, and should provide clues for future investigations into the roles of these neurons.

Cholinergic neurons in the motor ganglion have been implicated in the regulation of tail muscle contraction^21,32–34^. Here we show that one subtype of cholinergic neurons in the motor ganglion expresses *gnrh2* (Fig. 3A). These neurons extend axons posteriorly, but it is unclear whether they are motor neurons that directly innervate muscle cells or interneurons that connect to other CNS neurons in the caudal nerve cord. In the nerve cord, another class of cholinergic cells also expresses *gnrh2* (Fig. 3B). One possible role of cholinergic/GnRH neurons in the motor ganglion and the nerve cord may be the control of swimming behavior. These neurons may also play a role in metamorphosis, because GnRH has been suggested to be involved in the regulation of metamorphosis^13^.

Calcium imaging has been applied to studies of *Ciona* development^35–37^. These previous studies focused on Ca^2+^ transients in embryos but not in mature larvae. The present study for the first time reported the spatio-temporal patterns of Ca^2+^ transients in mature larvae of *Ciona*. Our observations included four novel findings: i) active Ca^2+^ transients in the proto-placode-derived aATENs, ii) periodic spikes in the motor ganglion of young larvae, iii) a correlation between Ca^2+^ transients and tail movements, and iv) active Ca^2+^ transients in ependymal cells of the nerve cord.

The proto-placode-derived aATENs share morphological and molecular properties with vertebrate olfactory neurons and are thought to be chemosensory cells^19^. However, olfactory receptors have not been identified in *Ciona*, and the chemical cues that stimulate aATENs are not known. Calcium imaging with *gnrh2*>G-CaMP8 could help us search for chemical cues that trigger the activation of aATENs in future studies.

Periodic Ca^2+^ transients observed in the motor ganglion of young larvae are reminiscent of the spontaneous rhythmic activities observed in the developing nervous systems of vertebrates^38–43^. These periodic neuronal activities are thought to be important for the development of neural circuits in the CNS and the retina^41,44,45^. Similar rhythmic oscillation of Ca^2+^ transients was reported in the developing motor ganglion of the *Ciona* embryo^37^. By contrast, we observed rhythmic Ca^2+^ transients in young larvae. The swimming behavior of *Ciona* larvae reveals ontogenic changes, and their photo-responsiveness appears a few hours after hatching^46–48^. Thus, the spontaneous rhythmic Ca^2+^ transients may play an important role in the neural circuit development of *Ciona* larvae.

We observed a correlation between the tail movements and Ca^2+^ transients in the motor ganglion and the nerve cord. This suggests that *gnrh2*-expressing cells are involved in the control of swimming locomotion. Ca^2+^ transients appeared when the tail stopped moving, and the Ca^2+^ signal was low when the tail was moving (Fig. 7B). In other words, the tail movement precedes the Ca^2+^ spike. This pattern suggests that these *gnrh2*-expressing cells are not motor neurons. In fact, the majority of the cells expressing *gnrh2* in the nerve cord are ependymal cells, and we observed Ca^2+^ transients in correlation with tail movements in ependymal cells. An intriguing possibility is that Ca^2+^ spikes are induced in ependymal cells by muscle contraction or motor axon excitation. If so, the ependymal cells may monitor the activity of muscle or motor neurons. It has been reported that various types of glia cells exhibit Ca^2+^ transients in response to neuronal activities and regulate neuronal functions in vertebrates^49–52^. The ependymal cells of *Ciona* larva may have similar regulatory roles, suggesting a deep evolutionary conservation of glia function between tunicates and vertebrates. Given the simplicity of its nervous system, the *Ciona* larva could serve as a unique model for the study of glia-neuron interaction.

In conclusion, the present study revealed the presence of dynamic Ca^2+^ transients of *gnrh2*-expressing cells at various sites in the *Ciona* larva. Our findings suggest a connection between the activity of *gnrh2*-expressing cells and the tail movements of the larva. An important yet unsolved question is whether GnRH2 is involved in these processes. Future studies should address the developmental and physiological roles of *gnrh2*-expressing cells and GnRH peptides based on the findings of this study.

## Methods

### Animals and embryos

Mature adults of *Ciona intestinalis* Type A (also called *Ciona robusta*) were provided by the Maizuru Fisheries Research Station of Kyoto University and by the Misaki Marine Biological Station of the University of Tokyo through the National Bio-Resource Project of the Ministry of Education, Culture, Sports, Science and Technology of Japan (MEXT), and were maintained in indoor tanks of artificial seawater (ASW) (Marine Art BR; Tomita Pharmaceutical, Tokushima, Japan) at 18°C. The adults were also collected from the pond on the Fukae campus of Kobe University, Kobe, Japan and from the fishing harbor in Murotsu, Hyogo, Japan. Eggs and sperm were obtained surgically from the gonoducts, and the eggs were fertilized *in vitro*. After insemination, the embryos were raised in ASW containing 50 μg/ml streptomycin sulfate (S6501; Sigma-Aldrich, St. Louis, MO, USA) at 18°C.

### Preparation of reporter constructs and electroporation

Construction of the *vglut*>*kaede* was described previously^53,27^. The *vacht*>*cfp* plasmid was made by inserting the 3.8-kb upstream region of *Ciona vacht*^25^ into the *Sal*I/*Bam*HI site of pSP-CFP^51^. The 2.4-kb upstream region of *Ciona fer2* (Gene ID KH.L116.39) was previously cloned into the pSP-CFP vector^27^. The reporter sequence was replaced with a DNA fragment coding for mCherry to generate *fer2>mcherry* using *Not*I/*Eco*RI sites. The *gnrh2*>*kaede* and *gnrh2*>*mcherry* plasmids were made by inserting the 4.3-kb upstream region of *Ciona gnrh2*^12^ into the *Xho*I/*Not*I sites of the pSP-Kaede vector and pSP-mCherry vector, respectively^54^. The *gnrh2* upstream region was also used to generate the *gnrh2*>*g-camp8* construct. The Kaede coding sequence of pSP-Kaede was replaced with a DNA fragment coding for G-CaMP8^31^ using *Not*I/*Eco*RI sites. The *gnrh2* upstream region was amplified from the *gnrh2*>*kaede* plasmid using a pair of nucleotide primers (5’-GAATCGGCCAACGCGGGATCCAGGAGCAGACGTCATAAGTA-3’ and 5’-TGACGCGGCCGCTGTTACGTTATCTCTCTAGAAG-3’), digested with *Bam*HI and *Not*I, and then inserted into the *Bam*HI/*Not*I sites upstream of the G-CaMP8 in the pSP vector. Plasmid DNA constructs were electroporated into fertilized *Ciona* eggs as described by Corbo et al.^55^.

### Immunofluorescent staining

Immunofluorescent staining was carried out according to the method described by Nishitsuji et al.^22^. Fluorescent images were obtained by using a laser scanning confocal microscope (FV1200 IX83; Olympus, Tokyo).

To visualize the localization of cell type-specific proteins, a mouse antiserum against *Ciona* VIAAT^21^ or a rabbit antiserum against *Ciona* CRALBP^23^ was diluted 1:1000 in 10% goat serum in T-PBS (0.1% Triton X-100 in PBS) and used as the primary antibody. The secondary antibody was an Alexa Fluor 594-conjugated anti-mouse IgG (A11005; Thermo Fisher Scientific) or an Alexa Fluor 594-conjugated anti-rabbit IgG (A11012; Thermo Fisher Scientific).

The primary antibodies used to visualize the localization of fluorescent reporter proteins were rabbit anti-Kaede polyclonal (PM012; Medical & Biological Laboratories, Nagoya, Japan; for Kaede), rabbit anti-green fluorescent protein (GFP) polyclonal (A11122; Thermo Fisher Scientific; for G-CaMP8 and CFP), rat anti-red fluorescent protein (RFP) monoclonal (5F8; ChromoTek GmbH, Martinsried, Germany; for mCherry), and rat anti-GFP monoclonal (GF090R; Nacalai Tesque, Kyoto, Japan; for G-CaMP8 double-stained with anti-CRALBP) antibodies. All the primary antibodies were diluted 1000-fold as described above. The secondary antibodies were an Alexa Fluor 488–conjugated anti-rabbit IgG (A11008; Thermo Fisher Scientific) for G-CaMP8 and CFP, an Alexa Fluor 488–conjugated anti-rat IgG (A11006; Thermo Fisher Scientific) for G-CaMP8, and an Alexa Fluor 594–conjugated antirat IgG (A11007; Thermo Fisher Scientific) for mCherry.

### *In vivo* Ca^2+^ imaging

Electroporated *Ciona* larvae expressing the G-CaMP8 transgene were placed in ASW on a 35-mm glass based dish (coverslip diameter 12 mm, #3931-035; Iwaki, Japan). For imaging, a microscope (IX81; Olympus, Tokyo) equipped with an electron multiplying charge-coupled device (EMCCD) camera (EVOLVE512; Photometrics, Tucson, AZ) was used. Images were taken with a 50-ms exposure time per 1 second and 1×1 binning. For each animal, 300 images were taken in 6 minutes. Changes in intracellular calcium concentrations were measured as the changes in the green fluorescence of G-CaMP8. The fluorescence intensity was spatially averaged in each region of interest (ROI). The fluorescence change was defined as *ΔF/F=(Ft – F0)/F0*, where *Ft* is the fluorescence intensity at time t, and *F0* is the baseline averaged for 4-5 s. The fluorescence change (Δ*F*/*F*) was calculated after subtracting the background fluorescence. The MetaMorph image analysis software system (Molecular Devices) was used for the analysis of the imaging. Image processing was also performed with ImageJ (U. S. National Institutes of Health, Bethesda, MD, USA, https://imagej.nih.gov/ij/).

## Acknowledgments

We thank the National BioResource Project of MEXT and all members of the Maizuru Fisheries Research Station and Yutaka Satou Lab of Kyoto University and the Misaki Marine Biological Station of the University of Tokyo for providing us with *C. intestinalis* adults. We also thank the Graduate School of Maritime Sciences, Kobe University for generously allowing us to collect *C. intestinalis* on the campus. This study was supported in part by Grants-in-Aid for Scientific Research from JSPS to T.G.K. (25650118, 25290067, 16H04724, 19H03213), J.N. (15H05723), and T.H. (16K07433, 19H03204). This study was also supported in part by research grants from the Takeda Science Foundation and the Hirao Taro Foundation of the Konan University Association for Academic Research.

---

**Movie S1.** Serial fluorescence images showing Ca^2+^ transients in the larva shown in Figure 4.

**Movie S2.** Serial fluorescence images showing Ca^2+^ transients in the larva shown in Figure 5.

**Movie S3.** Serial fluorescence images showing Ca^2+^ transients in the larva shown in Figure 6.

**Movie S4.** Serial fluorescence images showing Ca^2+^ transients in the larva shown in Figure 7.

## Notes

#### Summary of Updates

Abstract revised; main text revised; Figure 5 revised; Figure 6 revised; Figure 7 revised.

